# Climate change promotes hybridisation between deeply divergent species

**DOI:** 10.1101/066258

**Authors:** Daniele Canestrelli, Roberta Bisconti, Andrea Chiocchio, Luigi Maiorano, Mauro Zampiglia, Giuseppe Nascetti

**Author notes:** Correspondence author: Roberta Bisconti.

## Abstract

Rare hybridisations between deeply divergent animal species have been reported for decades in a wide range of taxa, but have often remained unexplained, mainly considered chance events and reported as anecdotal. Here, we combine field observations with long-term data concerning natural hybridizations, climate, land-use, and field-validated species distribution models for two deeply divergent and naturally sympatric toad species in Europe *(Bufo bufo* and *Bufotes viridis* species groups). We show that climate warming and seasonal extreme temperatures are conspiring to set the scene for these maladaptive hybridisations, by differentially affecting life-history traits of both species. Our results identify and provide evidence of an ultimate cause for such events, and reveal that the potential influence of climate change on interspecific hybridisations goes far beyond closely related species. Furthermore, climate projections suggest that the chances for these events will steadily increase in the near future.

## Introduction

Hybridisation is a widespread phenomenon in nature (Mallet, 2005). However, its frequency, diversity of outcomes, underlying mechanisms, its role in the evolutionary process, and how to deal with it in conservation biology have been controversial topics for more than a century (Arnold, 2006; Schwenk et al., 2008). Much of our knowledge about the link between hybridisation dynamics in animals and climate changes, comes from studies of hybrid zones (Hewitt, 2011), where the reshuffling of species’ ranges in response to changing climates brought into contact closely related and previously allopatric species. Pre-mating reproductive barriers could be incomplete between these species, and their genomes could still be porous to introgression, with several far reaching implications (Mallet, 2005; Arnold, 2006; Schwenk et al., 2008; Hewitt, 2011). Not surprisingly, species of ancient divergence and with a long-lasting history of sympatry have contributed the least to this body of knowledge (Mallet, 2005; Schwenk et al., 2008). These species have had ample opportunity to evolve strong pre-mating reproductive barriers, either as a by-product of a longer allopatric divergence or because of character displacement in response to natural selection (Coyne & Orr, 2004; Pfennig & Pfennig, 2009). Consequently, hybridisation events are extremely improbable between these species (e.g. Proietti et al., 2014), and their observation incidental in the wild.

We witnessed to one such event in southern Italy (on May 10^th^ 2014) between two toad species, the common toad *Bufo bufo* and the green toad *Bufotes balearicus* (Figure 1). They belong to the *Bufo bufo* and *Bufotes viridis* species groups, whose divergence has been estimated to the Oligocene (around 20-30 million years ago; Maxson, 1981; Garcia-Porta et al., 2012), and which have largely overlapping distributions in central, eastern, and southern Europe (Sillero et al., 2014). Although syntopy is not uncommon, especially in lowland areas, they show distinct spatio-temporal patterns of habitat use (reviewed in Lanza et al., 2006), making hybridisation at least three-times unexpected. First, they show markedly different breeding phenologies: *Bufo bufo* is an early and explosive breeder, while *Bufotes balearicus* is a late and prolonged breeder (Lanza et al., 2006). In Italy, breeding activities of *Bufo bufo* begin earlier in the year (early winter to early spring, with variation among sites at different altitude and latitude) and usually last 1-2 weeks. *Bufotes balearicus* starts breeding later (middle to late spring), and this activity may last 2-3 months. Furthermore, breeding activities in syntopic areas have been systematically reported as asynchronous, regardless of intraspecific differences between sites (Lanza et al., 2006). Second, *Bufo bufo* and *Bufotes balearicus* display differences in their altitudinal distribution. *Bufo bufo* breeding sites commonly occur from 0 to 2000 m above sea level (MASL), while *Bufotes balearicus* shows marked preferences for sites in lowland areas, rarely being observed above 1000 MASL (Romano et al., 2003; Spilinga et al., 2007). Third, in spite of the largely polytopic habits of these two species, differences exist in habitat and breeding site preferences. *Bufotes balearicus* favours open areas and bushlands and breeds in temporary shallow waters, and *Bufo bufo* most commonly inhabits forested habitats while using slow running or deeper, wider standing waters as breeding sites.

**Figure 1.**
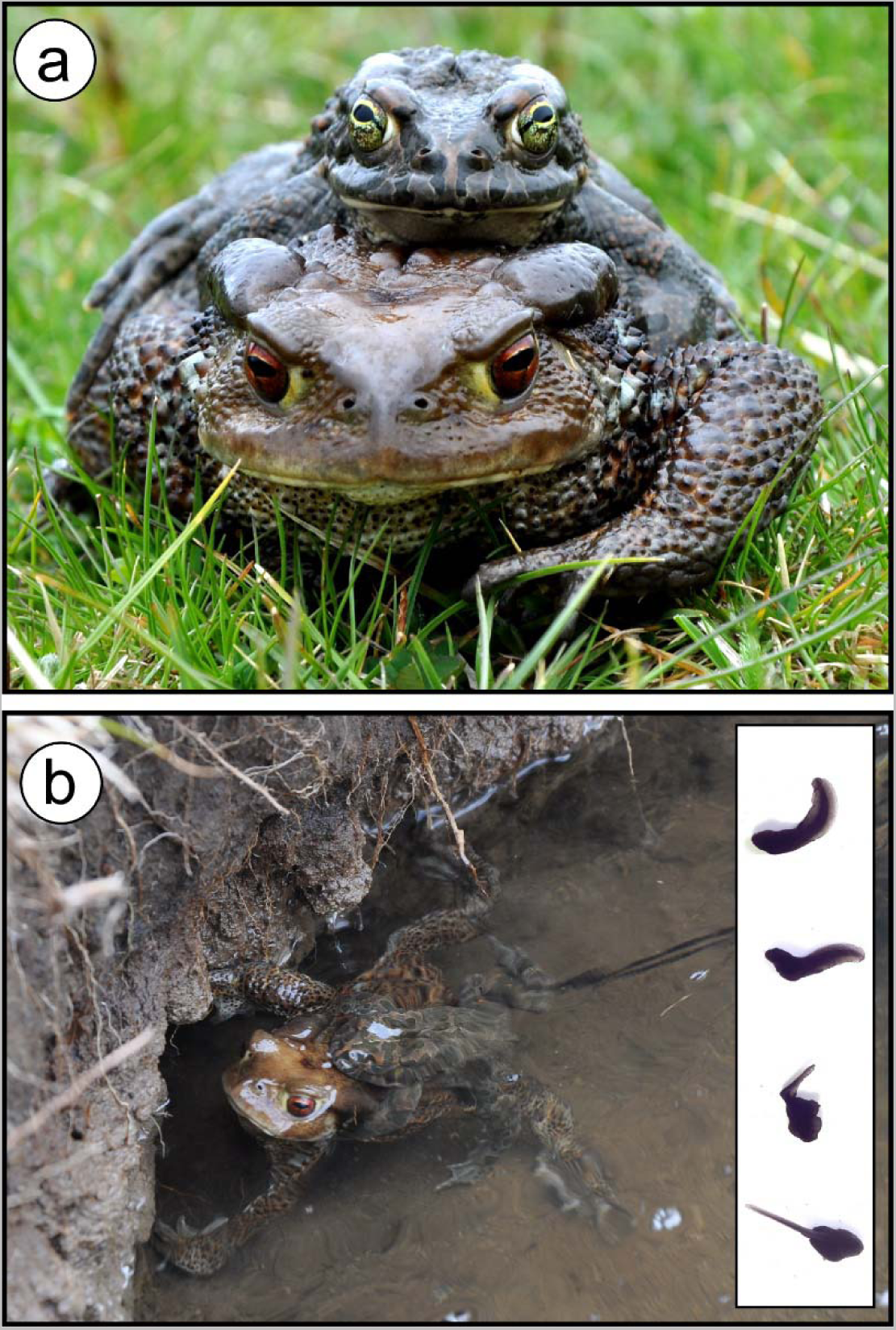
Interspecific hybridisation between the common toad (*Bufo bufo*) and the green toad (*Bufotes balearicus*) in the wild. The hybrid pair (a) was found spawning (b) on 10 May 2014, at lake Campo Maggiore, a high-elevation pond within the Partenio Regional Park, southern Italy (latitude: 40.9429° N; longitude: 14.7096° E; altitude: 1330 MASL). The majority of tadpoles from the hybrid egg-string reared under standard laboratory conditions were heavily malformed (inset), and none survived until metamorphosis; this pattern was not observed for control tadpoles from con-specific matings. (Photos: M. Zampiglia)

Remarkably, based on previous assessments (Carpino & Capasso, 2008), all these differences applied to toad populations at our study site as well. This site is a high-altitude pond located at the margins of a forested area (latitude: 40.9429° N; longitude: 14.7096° E; altitude: 1330 MASL; Figure 2). *Bufo bufo* was reported to breed at this site on February, whereas *Bufotes balearicus* was absent here and in neighbouring areas above 800 MASL at least until the year 2007, the year of the last herpetological assessment (Carpino & Capasso, 2008).

**Figure 2.**
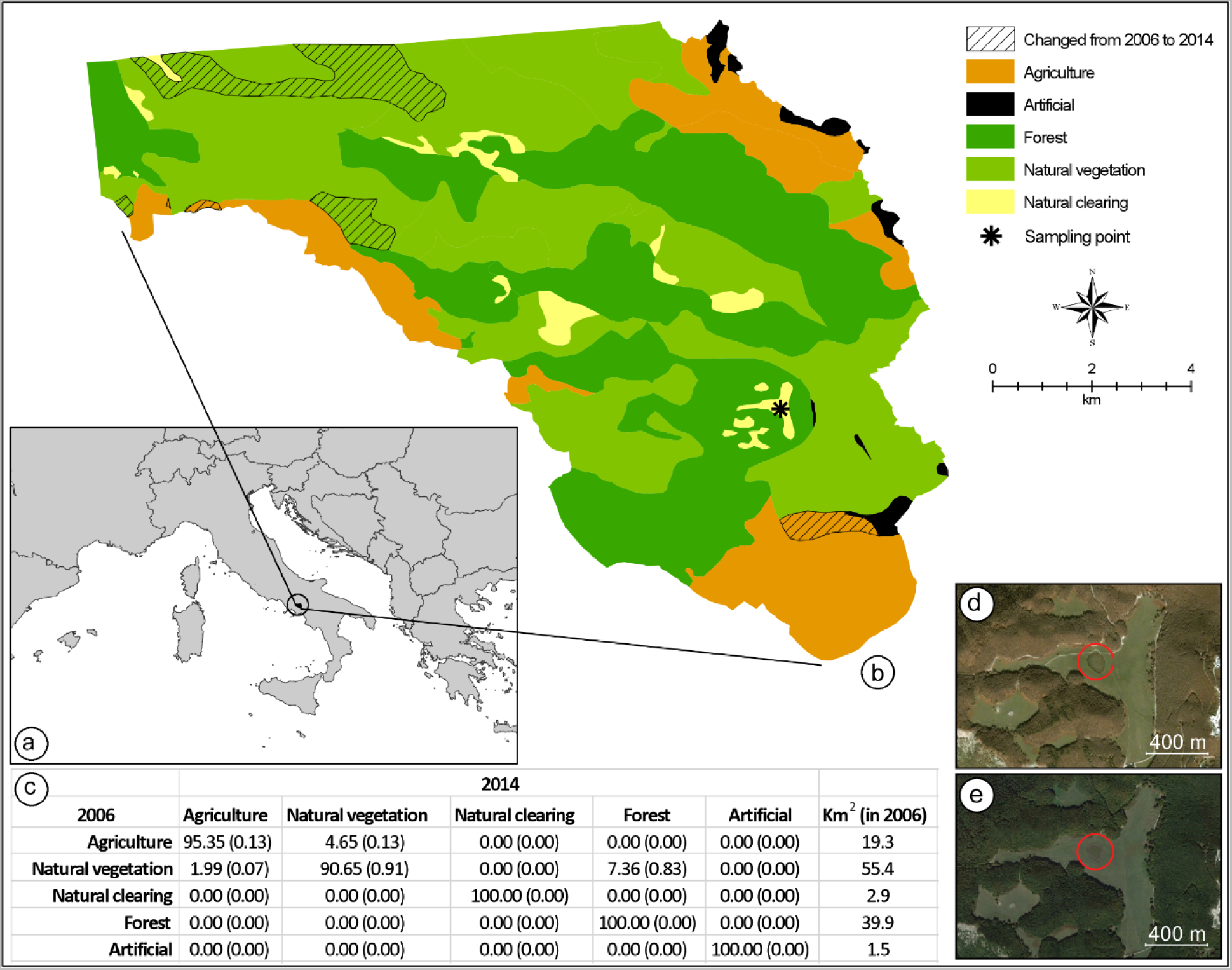
Land-use change detection analysis. (a) Location of the study area in Italy. (b) map of the land use in 2006, as obtained from direct interpretation of an aerial photo collected on 31 October 2006; in the same map, the exact location where the hybridisation event has been registered is indicated, as well as all areas where land use was different when compared to a second aerial photo collected on 9 October 2014. Both aerial photos were obtained from Google Earth Pro 7.1.5.1557 (Google Inc., Mountain View, California). (c) Average percent change (range of percent change in parenthesis) in land use classes from 2006 to 2014; total km^2^ area for each land cover class in 2006 is provided in the last column. Aerial photos of the breeding site and its neighbourhoods collected on 31 October 2006 (d), and 9 October 2014 (e).

Here, we combine our field observation with data from previous reports of similar hybridisation events for the *Bufo bufo* and *Bufotes viridis* groups in Europe along the last century, in order to study the causation of these ‘improbable’ events To this end, we examined the contribution of multiple factors, including all those commonly invoked to explain novel interspecific hybridisations among animal species in the wild.

## Materials & Methods

### Assessing natural hybridisation

We sampled and carried to the laboratory a strip approximately 1.5 m in length from the clutch laid by the hybrid pair on May 10^th^ 2014. We visually searched for additional clues of hybridisation at the breeding pond for the subsequent ten days. Although no further heterospecific pairs were observed, we collected and carried to the laboratory two additional egg-strings newly laid by unobserved parents. Fieldwork was approved by the Italian Ministry of Environment (permission number: 0042634 dated 7 August 2013).

In order to confirm the hybrid nature of the egg-string laid by the heterospecific pair (against the hypotheses of unfertilised eggs and of undetected homospecific paternity) and to address the parental species of the other egg-strings, we monitored egg and tadpole development under laboratory conditions and analysed the pattern of variation of individual larvae at diagnostic genetic markers. Tadpoles were reared under standardised light and food conditions, in plastic boxes (0.8 × 0.5 × 0.2 m) filled with oxygenated tap water. Larval mortality was checked twice daily, from hatching to metamorphosis.

Tadpoles of *Bufo bufo* and *Bufotes balearicus* can be distinguished by larval morphology (Ambrogio & Mezzandri, 2014) while hybrid tadpoles are usually heavily malformed (Montalenti, 1932; Montalenti, 1933). However, in order to achieve correct identification and to verify the absence of backcrosses between hybrids and parental individuals, we analysed genetic variation at the following allozyme loci: malate dehydrogenase (Mdh-1 and Mdh-2; EC 1.1.1.37), isocitrate dehydrogenase (Icdh-1 and Icdh-2; EC 1.1.1.42), and malate dehydrogenase NADP+-dependent (Mdhp-1; EC 1.1.1.40). Fifty tadpoles from each egg-string were euthanised using a 200 mg/L solution of MS222, 10 days after hatching, and stored at −80°C until subsequent analyses. The diagnostic value of each allozyme locus was verified through preliminary analyses of 20 individuals per species, sampled from two sites in neighbouring areas, where no evidence of potential hybridisation had been observed (*Bufo bufo*: 41.1737° N, 14.5834° E; *Bufotes balearicus*: 40.8866° N, 14.9318° E). Standard horizontal starch gel electrophoresis and zymogram visualisation were carried out, following previously published standard protocols (Harris & Hopkinson, 1976).

### Quantifying anthropogenic habitat change

In order to assess anthropogenic habitat change as a possible explanation for the hybridisation event we registered, we performed a land use change analysis considering an area of 119 km^2^, inside the Partenio Regional Park (Regione Campania, Italy). The clearing where the hybridisation event was registered is located in the middle of the study area, approximately 15.6 km from the northern boundary and 2 km from the south-eastern boundary (Figure 2). We obtained a map of the area from Google Earth Pro 7.1.5.1557 (Google Inc., Mountain View, California). By using the “historical imagery” tool, and keeping the extent and resolution of the map constant, we selected two images: one from 10 October 2014, five months after the hybridisation event, and one from 31 October 2006, a few months before the last assessment that confirmed the absence of *Bufotes balearicus* from the site (Carpino & Capasso, 2008). The two images were imported into ArcGIS 10.3.1 (ESRI ©), with a resolution of 4.5 m per pixel, and were georeferenced using administrative boundaries as reference points (RMS error = 6.23 m for the 2014 image; RMS error = 4.48 m for the 2006 image). The images were interpreted using direct recognition (Campbell, 1978), considering five discrete land use classes that hold a clear ecological importance for both toad species: agriculture, forests, natural clearings, natural vegetation (other than forests), and artificial areas. For both images, a vector layer (format shapefile, ESRI ©) was produced at a 1:25,000 scale. To perform the land cover change analysis, following Falcucci et al. (2007), each vector layer was transformed into a raster layer using 4 different pixel resolutions: 25 m, 50 m, 75 m, and 100 m. The change detection analysis was performed for each pixel resolution, resulting in an average percentage change for every land cover class.

### Climate influence on altitudinal distribution pattern

Based on historical records of occurrence (Carpino & Capasso, 2008), and considering the species’ altitudinal distributions in peninsular Italy (Lanza et al., 2007; Guarino et al., 2012), the presence of a *Bufo bufo* population at the study site was expected, whereas the presence of *Bufotes balearicus* was not expected, either at this site or within neighbouring, high altitude areas. Therefore, we focused the following analyses on the latter.

To address the plausibility of climate forcing on recent altitudinal distribution changes for *Bufotes balearicus*, we calibrated a correlative species distribution model in peninsular Italy considering the presence of the species across the 21^st^ century. Then we projected the distribution model to the current climate (average over 2007-2013) and to the future (average 2070-2100). The model was calibrated considering six bioclimatic variables theoretically important for the presence of the species: temperature seasonality, mean temperature of the warmest quarter, mean temperature of the coldest quarter, temperature annual range, precipitation seasonality, and precipitation of the coldest quarter. We obtained all climate variables at 1 km resolution from WORLDCLIM (Hijmans et al., 2005), which provides climate layers representative of the 1950–2000 time frame. To obtain the corresponding climate variables for the 2007–2013 time frame, we followed the procedure presented in Maiorano et al. (2013) and considered monthly temperature and precipitation values (spatial resolution equal to 50 × 50 km) from the Climatic Research Unit of the University of East Anglia (database: CRU TS3.22; Harris et al., 2014). To downscale the CRU database to the 1 km^2^ of the WORLDCLIM database, we first calculated climate anomalies by contrasting monthly temperature and precipitation values for 2007–2013 against the 1950–2000 climate data, as obtained from the same CRU TS3.22 database. Anomalies were calculated as absolute temperature difference (Δ°C) and relative precipitation differences (% change). By using bilinear resampling, we downscaled the anomalies to 0.0083° of spatial resolution (≈ 1km). Then, in order to obtain monthly maps of temperature and precipitation for 2007–2013, we applied the anomaly corrections to the WORLDCLIM climate layers. Finally, we calculated all the derived climate maps mentioned above.

The bioclimatic layers for 2070–2100 were obtained directly at the resolution of 1 km^2^ from the WORLDCLIM database considering 3 emission scenarios (A1B, A2, and B1), and many different global circulation models (24 GCMs for the A1B emission scenario, 19 GCMs for A2, and 18 GCMs for B1) developed under the 4^th^ Assessment Report of the Intergovernmental Panel on Climate Change (IPCC, 2007).

To calibrate the models, we used the ensemble forecasting approach (Araújo & New, 2007) implemented in BIOMOD, a bioclimatic niche modelling package for the R environment (Thuiller et al., 2009). We used the following eight models: (i) generalised linear models, (ii) generalised additive models, (iii) classification tree analysis, (iv) artificial neural networks, (v) generalised boosted models, (vi) random forests, (vii) flexible discriminant analysis, and (viii) multivariate adaptive regression spline. All models were calibrated over the entirety of peninsular Italy south of the Po river (212,460 km^2^), with 350 points of presence for *Bufotes balearicus* collected before 2000, plus 10,000 background points (see Supplemental Information S1). All models were evaluated using a repeated split-plot procedure (70% of the data used for calibration, 30% left apart for evaluation; the entire procedure repeated 10 times for each model; Thuiller et al., 2009), and by measuring the area under the receiver operating characteristic (ROC) curve (AUC) (Swets, 1988). All models with AUC values greater than 0.7 (Swets 1988) were projected over the entire study area using the 1950–2000, the 2007–2013, and the 2070-2100 climate layers. We measured the minimum probability of presence obtained in correspondence of the available points of presence for 1950–2000, and we used this threshold to define areas of species presence (all areas above this minimum threshold of probability) in all periods considered. Moreover, considering 100 m wide elevation classes, we calculated the elevation-specific average probability of presence for all three periods, and obtained a model of the probability of presence for *Bufotes balearicus* along the elevation range in peninsular Italy (Figure 3).

**Figure 3.**
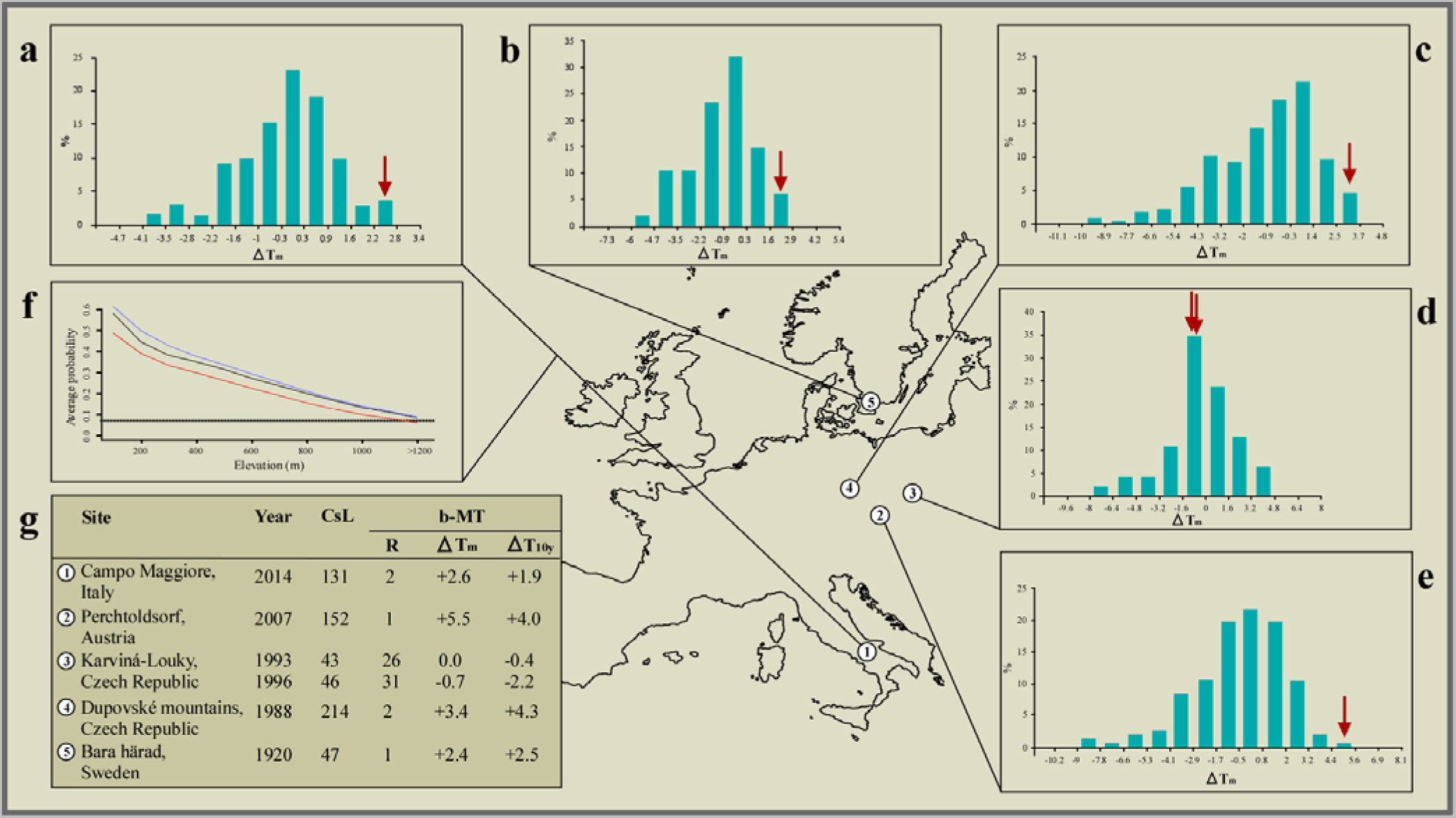
Climate correlates of the interspecific hybridisation events observed in the wild between species of the common toad *(Bufo bufo)* and the green toad *(Bufotes viridis)* species groups. Bar plots showing frequency distribution (percent) of bimonthly mean temperature deviations (ΔTm) from the 1961–1990 average, compared to the two months preceding the breeding activity at each geographic region: (a) December to January (Italy, this study), (b) February to March (Sweden, Lang, 1926), and (c-e) January to February (Czech Republic, Vlček, 1995, Vlček, 1997, Zavadil & Roth, 1997, respectively). Values for the years when hybrid mates were observed are marked using red arrows. Optimal bar width was computed for each climatic series following the Freedman–Diaconis rule. (f) Average probability of presence vs elevation at sea level (m) as modelled for the pre-2000 climate (red line), the 2007–2013 climate (solid black line), and the 2070-2010 climate (blue line); the black dotted line indicates the minimum plausible level of probability of presence, above which the species can be considered present, while below is considered absent. (g) Mean temperature data for each site, and year of observation of interspecific mates. CsL: climatic series length, in years, before the observed event; b-MT: bimonthly mean temperature; R: rank over the entire climatic series (1= mildest); ATm: deviation from the 1961–1990 average temperature (°C); ΔT10y: deviation from the preceding 10-year average temperature (°C).

We further investigated the plausibility of a link between climate change and altitudinal shifts by turning the model prediction into a working hypothesis. Based on the model results, we selected a geographic area close to our study site, and carried out field searches for further, unknown sites of occurrence of *Bufotes balearicus,* above 1200 MASL. To select the geographic area, we adopted the following criteria: (i) location on a mountain massif, as close as possible to our study site; (ii) presence of potential breeding sites (e.g., ponds) at altitudes ≥1200 MASL; (iii) presence of *Bufotes balearicus* populations at lower elevations along the same mountain; (iv) absence of obvious anthropogenic habitat discontinuities between low and high altitude areas. Accordingly, we identified the Picentini Mountains (within the Picentini Mountains Regional Park, roughly located 25 Km southeast of our study site) as an area of best fit for our criteria. Field searches began on 2 May 2015, and lasted until the first evidence of *Bufotes balearicus* in the area was found (21 May). The rationale underlying this experimental integration was as follows: although failure to identify new high altitude sites of occurrence would not be strong evidence against a role of climate change in promoting altitudinal shifts at lake Campo Maggiore, or elsewhere, a positive result would provide support for the model prediction, and therefore, support the hypothesis that our initial finding belongs to a suite of events promoted by climate change.

### Climate influence on breeding phenology

Our observation of the hybrid pair in May 2014 suggests delayed breeding activity of *Bufo bufo* causing an overlap with the normal breeding period of *Bufotes balearicus* (Lanza et al., 2006; Carpino & Capasso, 2008; Guarino et al., 2012). Therefore, subsequent analyses were focused on *Bufo bufo.* Notably, while there is strong evidence for a link between the breeding phenology of *Bufo bufo* and annual temperature cycles (Reading, 1998; Reading, 2003; Tryjanowski et al., 2003), the same does not hold true for species of the *Bufotes viridis* group (including *Bufotes balearicus).*

A search of academic and grey literature revealed five additional observations of hybrid pairs among representatives of the two species groups in four different locations (Figure 3): two sites located in the Czech Republic (Vlček, 1995; Vlček, 1997; Zavadil and Roth, 1997), one in Sweden (Lang, 1926), and one in Austria (Duda, 2008).

The annual activity cycle of *Bufo bufo* populations can be affected by several environmental features, including climate, and the five sites (including our observation) span a wide range of latitudes. Thus, rather than considering average winter temperatures, we based our analysis on the period when this species begins its breeding activity in each area, according to regional atlases and databases (Gilsen & Kauri, 1959; Cabela, Grillitsch & Tiedemann, 2001; Nečas et al., 1997; Guarino et al., 2012;). In addition, previous studies suggested that the beginning of this activity is linked to the average temperatures of the preceding 1–2 months (Reading, 1998). Therefore, in our testing for a link between hybridisation events and climate anomalies, we set the period of interest to the two months preceding the usual start of the breeding activity, for each geographic area. Accordingly, we analysed date ranges covering December to January for the site in southcentral Italy, January to February for the sites in Czech Republic and Austria, and February to March for the site in Sweden.

Long-term climate data for our study site were provided, by the Montevergine Observatory (40.9360° N; 14.7288° E), as monthly averages since the year 1884. In order to gain climate data for the four sites of past hybridisation, we searched the NOAA database (available at http://gis.ncdc.noaa.gov) of monthly observational data using the following two criteria: (i) climate station closest to the site of interest and (ii) time series of at least 40 years before the year of the observed hybridisation event. The following stations best matched these search criteria: Koebenhavn Landbohojskolen, Denmark (Id: DA000030380; Latitude: 55.683° N; Longitude: 12.533° E); Praha Klementinum, Czech Rep. (Id: EZE00100082; Latitude: 50.090° N; Longitude: 14.419° E); Wien, Austria (Id: AU000005901; Latitude: 48.233° N; Longitude: 16.35° E); OravskaLesna, (Id: LOE00116364; Latitude: 49.366° N; Longitude: 19.166° E).

For each climatic series retrieved, we analysed bimonthly average temperatures considering the entire temporal series, and the 10 years preceding the hybridisation event, i.e. a time-lapse approximating the average lifetime of a toad in the wild (Lanza et al., 2006).

To test the null hypothesis that an association between hybridisation events and temperature anomalies was due to chance alone, we carried out binomial probability tests. We set the probability threshold of a single event to 0.02, based on the highest value calculated for the ratio between year rank (mildest = 1st rank) and climatic series length (i.e. the first out of 47 available years from the climatic station DA000030380). Since hybridisation events were both spatially and temporally distant, data independence was assumed. However, to err on the side of caution, we carried out the analyses considering the two observations in eastern Czech Republic, as both independent and fully dependent (i.e. as a single observation); then we took the highest value as the confidence level for accepting/rejecting the null hypothesis stated above.

Finally, the paucity of hybridisation events recorded qualifies these events as rare, and testifies to the strength of the pre-mating isolation mechanisms. On the other hand, given such rareness, we cannot exclude the occurrence of potentially unobserved, unreported, or undetected hybridisation events. Thus, we explored how potentially unknown events could affect the significance of our test. To this aim, we carried out additional binomial probability tests by progressively increasing the number of events while leaving the number of ‘successes’ unchanged. The null hypotheses of no association was rejected at the nominal probability threshold α = 0.05.

## Results and Discussion

At the time of our observation (May 10^th^ 2014), we counted 9 males, 3 females and 8 juveniles (22-26 mm long; presumably 1 year old) of *Bufotes balearicus,* plus 2 female *Bufo bufo,* and various newly spawned egg-strings.

All tadpoles from the putatively hybrid egg-string were identified as first-generation hybrids by their heterozygote status at all loci analysed. In line with previous findings (Montalenti, 1932; Montalenti, 1933), most of them were heavily malformed (see Figure 1), and none reached the metamorphosis. Instead, at all the loci analysed, tadpoles from the additional two clutches sampled at the breeding site were homozygotic for *Bufotes balearicus* diagnostic alleles, and were thus identified as belonging to this parental species. As expected, they did not show abnormalities, neither in the external morphology nor in the ontogenetic pathway.

Despite their wide sympatry, ease of observation, and more than a century-old knowledge of hybridisation in laboratory crosses, our literature searches for previous reports of interspecific breeding pairs in the wild, identified just 5 additional observations within a 94-year time span (Sweden, Lang, 1926; Czech Republic, Vlček, 1995, Vlček, 1997, Zavadil & Roth, 1997; Austria, Duda, 2008).

Three main hypotheses have been invoked to explain recently established interspecific hybridisations among animal species, and may have played a role in the present case by promoting syntopy and breeding season overlap (Crispo et al., 2011; Chunco, 2014): species translocations, anthropogenic habitat degradation (a derivation of the Anderson’s ‘hybridisation of the habitat’ model; Anderson, 1948) and climate changes.

In the case of *Bufo bufo* and *Bufotes viridis,* a species translocation can be firmly excluded in all the reported cases, based on the extensive knowledge of their natural geographic distributions (Lanza et al., 2006; Sillero et al., 2014), as well as on the fossil data of both species in Europe (Martin & Sanchiz, 2011).

Anthropogenic habitat degradation has been proposed as a main causative agent in some case (Duda, 2008). By reducing the diversity and number of potential breeding sites in a given area, physical alterations of habitat could promote syntopy of previously allotopic populations. Although plausibly contributing, this hypothesis cannot explain the entire pattern, and it does not apply to all cases. Our study site (but see also Zavadil & Roth, 1997) is located within a protected area established in 1993, and an analysis of contemporary and historical aerial photos of this site and neighbouring areas clearly show the absence of any physical alterations of potential relevance for the two species (Figure 2). Moreover, habitat degradation could not explain the overlap of the two breeding seasons. Climate changes, however, significantly improve our ability to explain the occurrence of hybridisation events between these species.

By promoting a recent altitudinal migration of *Bufotes balearicus* from neighbouring, lower altitude sites, the ongoing climate warming engendered the unexpected syntopy at our study site. Support to this argument (the only alternative to recent translocation), comes from our models of the distribution of *Bufotes balearicus* in peninsular Italy, based on a set of known occurrences collected before year 2000, and projected to the average climate over the period 2007-2013. Indeed, our models indicated that the species’ presence above 1200 MASL was highly improbable under pre-2000 climate, but became plausible during 2007-2013 (Figure 3). Furthermore, projecting the models under future climate projections for the time period 2070-2100 under different emission scenarios the general pattern remains unchanged (Figure 3). The reliability of the models was clearly confirmed by the field-validation procedure (see Methods).

Indeed, our field searches of *Bufotes balearicus* at high-altitude sites of predicted presence in post-2000 projections were successful. We found a previously unreported site of occurrence within the Picentini Mountains (Latitude: 40.8251°N; Longitude: 14.9864°E; roughly 25 Km south-east of the study site), thus confirming that upward migrations of *Bufotes balearicus* are ongoing, as predicted by our bioclimatic model (see also Zavadil & Roth, 1997).

Besides being a co-factor for syntopy, climate changes also contributed to the hybridisation events by promoting an overlap of the breeding activities. Analysing long-term climate series, we found that the years when hybridisation events were recorded in Europe (including our observation) ranked 1st or 2nd hottest on record at most site, over time series from 47 to 214 years-long. Moreover, bimonthly mean temperatures at these sites were 2.4 °C to 5.5 °C above the 1961-1990 averages, and 1.9 °C to 4.3 °C above the preceding 10-years averages (Figure 3). Binomial probability tests allowed us to reject the null hypothesis of random association between hybridisation events and extremely mild winters, with very high confidence (binomial probability: P ≤ 2.3×10^6^). Also, additional binomial probability tests, carried out in order to explore how unrecorded events could affect the significance of our test, indicated that the null hypothesis of random association was rejected (at *α* = 0.05) until the number of events was ≥ 77, while leaving unchanged the number of ‘successes’.

Although *Bufo bufo* is expected to bring forward its breeding activity after mild winters (Reading, 1998; Reading, 2003; Tryjanowski et al., 2003), at least three lines of support may help explaining this apparently counterintuitive pattern. In years when the breeding season begins earlier (after a mild winter), breeding has been observed to last longer (Gittins et al., 1980). Furthermore, a second and lower peak of breeding activity has been often observed later in the season (Pages, 1984; Reading, 1998), especially after mild winters (Reading, 1998). Finally, extensive ecophysiological investigations on bufonid toads, including *Bufo bufo,* indicate that increased temperatures during hibernation lead to significant alterations of several processes affecting the breeding activity, including body size condition, annual ovarian cycle, and seasonal synchronisation of breeding (Jørgensen, 1992).

Our analyses do not indicate climate change as the single explanatory factor. The environmental contexts in which interspecific interactions occur, the diverse forms of habitat disturbance, or behavioural changes during altitudinal migrations (Canestrelli et al., 2016a; Canestrelli et al., 2016b), might be locally influential. Nonetheless, these analyses clearly show that climate changes played a fundamental part in promoting hybridisation events. In light of the direction of these changes (IPCC, 2014), and of the results of our modelling exercise, we hypothesise that these events will become progressively more common in the near future. Most importantly, our results reveal a wider potential influence of climate changes on interspecific reproductive interactions, particularly in the many instances where climate-driven asynchrony and/or allotopy are integral components of the reproductive isolating barriers.

Hybridisation events among non-closely related species are generally believed to yield events that are transient, and potentially affecting local population demography at most, because strongly maladaptive (Rhymer & Simberloff, 1996; Malone & Fontnot, 2008). Nevertheless, there may be exceptions, whereby the effects of maladaptive processes propagate from population to community level (Farkas, 2015). Moreover, as revealed by years of investigation on the hybridisation process in several animal taxa, including amphibians, new evolutionary pathways have been sometime opened by such rare and maladaptive events (Arnold, 2006).

## Acknowledgments

We thank Maurizio Severini and Graziano Crasta for statistical advice, Vincenzo Capozzi for providing climatic data for the Montevergine Observatory, and Paola Arduino for providing support during laboratory procedures.

